# Dynamic recruitment of single RNAs to processing bodies depends on RNA functionality

**DOI:** 10.1101/375295

**Authors:** Sethuramasundaram Pitchiaya, Marcio D.A. Mourao, Ameya Jalihal, Lanbo Xiao, Xia Jiang, Arul M. Chinnaiyan, Santiago Schnell, Nils G. Walter

**Author notes:** Correspondence (S.P.) and (N.G.W.).

## Abstract

Cellular RNAs often colocalize with cytoplasmic, membrane-less ribonucleoprotein (RNP) granules enriched for RNA processing enzymes, termed processing bodies (PBs). Here, we track the dynamic localization of individual miRNAs, mRNAs and long non-coding RNAs (lncRNAs) to PBs using intracellular single-molecule fluorescence microscopy. We find that unused miRNAs stably bind to PBs, whereas functional miRNAs, repressed mRNAs and lncRNAs both transiently and stably localize within either the core or periphery of PBs, albeit to different extents. Consequently, translation potential and positioning of *cis*-regulatory elements significantly impact PB-localization dynamics of mRNAs. Using computational modeling and supporting experimental approaches we show that phase separation into large PBs attenuates mRNA silencing, suggesting that physiological mRNA turnover predominantly occurs outside of PBs. Instead, our data support a role for PBs in sequestering unused miRNAs to regulate their surveillance and provides a framework for investigating the dynamic assembly of RNP granules by phase separation at single-molecule resolution.

## INTRODUCTION

Sub-cellular, membrane-free granules have emerged as critical components of normal biology and pathophysiology (Banani et al., 2017; Shin and Brangwynne, 2017), owing to their key role in spatial regulation of gene expression (Martin and Ephrussi, 2009; Spector, 2006). Processing bodies (PBs) are one such class of ribonucleoprotein (RNP) granules that persist during cellular homeostasis and are enriched for RNA processing and degradation enzymes (Eulalio et al., 2007a; Parker and Sheth, 2007). These granules are observed in almost all eukaryotes, ranging from yeast to mammals, and have been implicated in multiple biological processes, including oogenesis, progression through early development, and mediation of neuroplasticity (Buchan, 2014).

Mammalian PBs have been hypothesized to assemble via an RNA dependent phase separation model (Banani et al., 2017; Schutz et al., 2017; Teixeira et al., 2005), wherein multiple translationally repressed RNPs are concentrated within dense foci through strong multivalent interactions and individual or oligomeric RNPs loosely interact with these dense regions to create dynamic shells. While PBs display a wide array of dynamic behaviors (Aizer et al., 2008), the intra-granular RNP dynamics and RNP recruitment – processes that mediate the maintenance, maturation and putative gene regulatory functions of PBs – are largely unknown. Compositionally, PBs are predominantly composed of translationally repressed messenger RNAs (mRNAs) (Hubstenberger et al., 2017), but they also contain mRNA-regulating miRNAs (Liu et al., 2005) and, to a lesser extent, regulatory long non-coding RNAs (lncRNAs) (Hubstenberger et al., 2017). Yet, it is still unclear if the principles governing RNP-PB localization dynamics are similar or distinct for different classes of PB-localized RNAs.

PBs have been functionally associated with translational repression and/or degradation of mRNAs (Hubstenberger et al., 2017; Liu et al., 2005). In support of this functional connection, the inhibition of miRNA biogenesis or activity, while alleviating both translational repression and degradation of mRNAs, also causes the disappearance of PBs (Chu and Rana, 2006; Jakymiw et al., 2005; Pauley et al., 2006). By contrast, other reports have postulated that translationally repressed mRNAs are instead protected from degradation via accumulation in PBs (Buchan, 2014; Hubstenberger et al., 2017; Schutz et al., 2017). Consistent with these observations, the lack of microscopically visible decay fragments within PBs (Horvathova et al., 2017) and the persistence of gene repression in the absence of microscopically visible PBs (Eulalio et al., 2007b; Stalder and Muhlemann, 2009) support a role for PBs in mRNA storage. However, all of these studies rely on visualizing higher order PB structures as large microscopically visible foci (> 250 nm), whereas the potential for smaller, sub-microscopic PBs (as smaller protein complexes or individual PB components) contributing to RNA regulation has not been explored.

Here, we aim to address the following unresolved questions – 1) Are RNA-PB colocalizations dynamic? 2) Are all types of cellular RNAs, coding or non-coding, small or large, recruited to PBs in similar fashion? 3) What RNA-intrinsic factors contribute towards RNA-PB colocalization? 4) What functional effect does PB colocalization have on an RNA? To this end, we developed a methodology to simultaneously observe single molecules of RNA (miRNAs, mRNAs or lncRNAs) and individual PB foci inside both living and fixed human cells. We demonstrate that a majority of miRNAs and repressed mRNAs are stably anchored within the core of PBs, whereas translationally active mRNAs and lncRNAs associate with PBs only transiently at both the core and periphery of PBs, suggesting a strong correlation between PB-localization and RNA class. Furthermore, we find that unused (target-less) miRNAs are enriched at PBs and that the 3‘ versus 5‘terminal positioning of cis-regulatory miRNA response elements (MREs) dictates the PB localization patterns and dynamics of mRNAs. Finally, in silico modeling and experimental validation through hyperosmotic-stress induced phase separation suggests that the stochastic collision of mRNAs with freely diffusing, sub-microscopic PBs leads to more efficient mRNA regulation than their recruitment to microscopic PBs. Taken together, our observations reveal the fundamental principles governing the dynamic recruitment of different classes of cellular RNAs to PBs, a molecular basis for the assembly of membrane-less PB granules, and a novel potential function for PBs in accumulating target-less miRNAs for miRNA surveillance.

## RESULTS

### Super-resolved single-molecule fluorescence microscopy probes RNA-PB interactions

To dissect the localization dynamics of RNAs at PBs, we first created a U2-OS cell line that stably expressed GFP tagged Dcp1a, an mRNA decapping co-activator and PB marker (Aizer et al., 2008; Hubstenberger et al., 2017). We selected a clone (hereon termed UGD) with similar number and composition (based on colocalization with other PB markers) of Dcp1a foci as endogenously found in U2-OS cells (Figure S1A-D). Next, we chose two distinct methods for fluorescent labeling and intracellular delivery of RNAs based on their size. Mature regulatory miRNAs, whose size (~22 nt per strand) precludes endogenous labeling strategies (Pitchiaya et al., 2014), were chemically synthesized with a fluorescent Cy5 dye at the 3‘end of one of their two complementary strands, typically the guide strand. Since transfection results in the sequestration of RNA within subcellular vesicles (Cardarelli et al., 2016), we chose to deliver these miRNAs via microinjection (Figure 1A-C), which defines a clear starting point for our assays by instantaneously exposing RNAs to the cellular milieu (Pitchiaya et al., 2012; Pitchiaya et al., 2017; Pitchiaya et al., 2013). We confirmed that fluorophore labeling and microinjection did not affect the gene-repressive function (Figure S1E-H) of a tumor suppressive let-7 (l7/l7* and l7-Cy5/l7*) miRNA (Pitchiaya et al., 2012). Larger (> 200 nt) mRNAs and lncRNAs, which are difficult to synthesize chemically, were instead endogenously expressed and tagged via a modified version of the MS2-MCP labeling system (Fusco et al., 2003). Briefly, UGD cells were transfected with plasmids bearing the MS2-tagged gene of interest (GOI) and Halo-tagged MCP. Co-expression of these two constructs within the same cell results in Halo-MCP bound MS2-RNAs (Figure 1D-F), which are then visualized in live cells by coupling the Halo tag with the cell-permeable, fluorescent dye JF646 (Grimm et al., 2015). mRNAs and lncRNAs in fixed cells were visualized by single-molecule fluorescence *in situ* hybridization (smFISH) (Raj et al., 2008). We confirmed that an MS2-MCP tagged mRNA bearing the firefly luciferase (FL) coding sequence (CDS) and an artificial 3‘untranslated region (3‘UTR) bearing six tandem miRNA response elements (MREs) for let-7 (l7-6x) was translated and regulated much like its untagged counterpart (Figure S1I). We additionally showed that the tagged lncRNA THOR, which binds PB-enriched IGF2BP1 protein (Hubstenberger et al., 2017), mediated an oncogenic phenotype by promoting cell growth and stimulating oncogene expression (Figure S1J-L) as expected (Hosono et al., 2017). We then adapted a super-registration fluorescence microscopy-based tool (Grunwald and Singer, 2010) that measures intermolecular distances of spectrally distinct fluorescent molecules with nanometer (nm) precision, thereby enabling the capture of transient RNA-PB interactions in living cells and the precise quantification of RNA localization patterns within PBs in fixed cells (Methods). By combining this tool with intracellular single molecule, high-resolution localization and counting (iSHiRLoC) (Pitchiaya et al., 2012; Pitchiaya et al., 2017; Pitchiaya et al., 2013) and single mRNA imaging, we were able to probe the PB-associated dynamics, stoichiometry and localization of either microinjected miRNAs (Figure 1A-C, Supplementary movie 1) or ectopically expressed MS2-MCP tagged m/lncRNAs (Figure 1D-F) at a spatial accuracy of 30 nm and temporal resolution of 50 ms. We found that RNAs diffused ~100-1,000-fold slower while at PBs compared to the cytosol (Figure 1G), supporting the notion that they physically dock to form higher order complexes at PBs. Concordantly, cytoplasmic RNAs were predominantly monomeric, whereas a significant fraction of RNAs in PBs appeared multimeric (Figure 1H-J).

**Figure 1.**
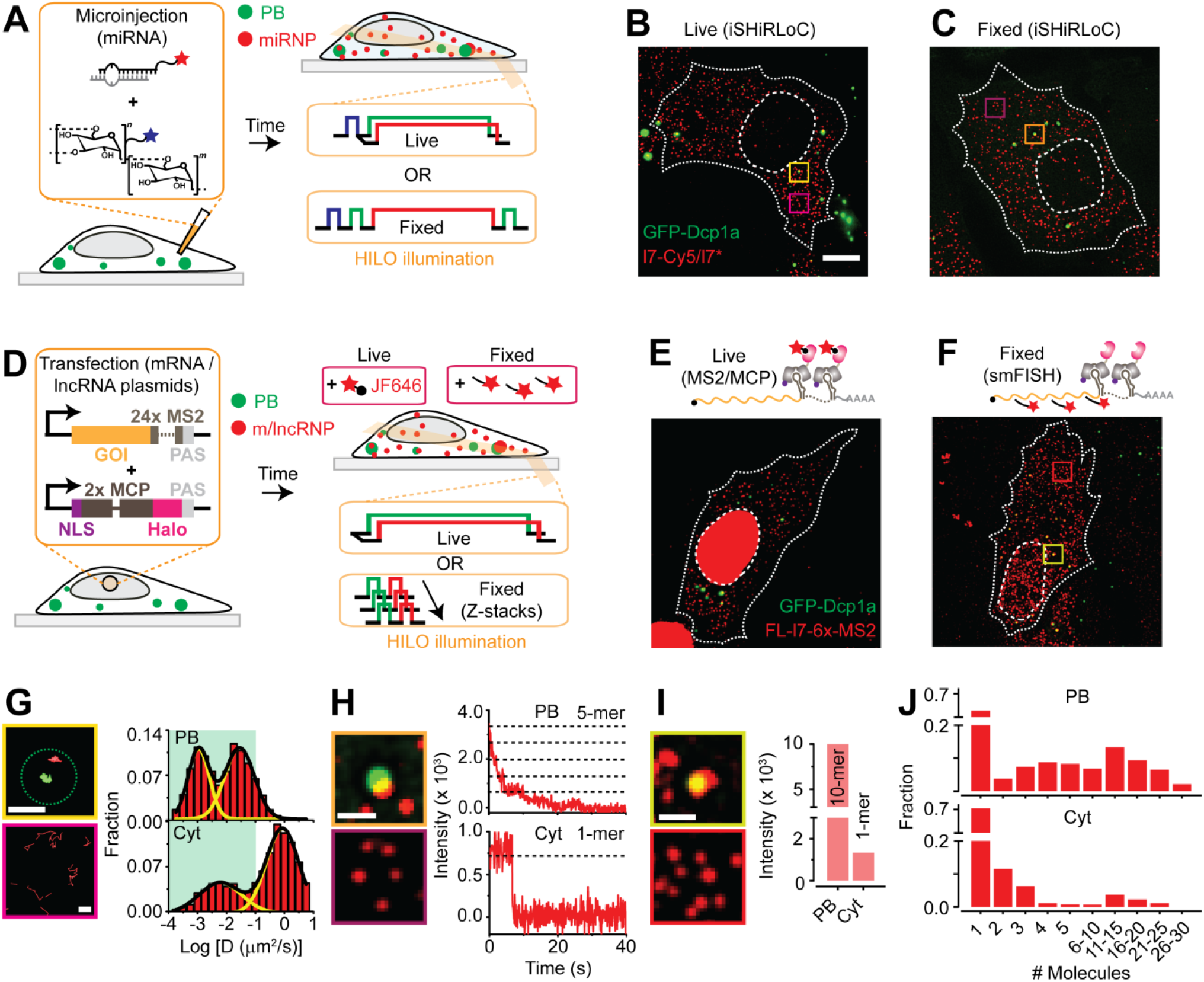
A super-resolution imaging tool for probing RNA-granule dynamics and colocalization. (A) Schematic of iSHiRLoC assay for probing miRNA-PB dynamics and colocalizations. (B and C) Representative pseudo-colored and contrast-adjusted images from live-cell imaging (B) and fixed cell imaging (C) assays of UGD cells expressing < GFP-labeled PBs (green) that were microinjected with l7-Cy5/l7* miRNA (red) and imaged 2 h post injection. Scale bar, 10 μm. (D) Schematic of assay for probing m/lncRNA-PB dynamics and colocalizations. NLS – nuclear localization signal, GOI – gene of interest and PAS – polyA signal. (E and F) Representative pseudo-colored and; contrast-adjusted images from live- and fixed-cell imaging assays respectively, of UGD cells (green) that were transfected with the FL-l7-6x-MS2-Halo-MCP constructs (red) (G) Representative single-particle trajectories of PBs (green) and l7-Cy5/l7* miRNA (red) from yellow and magenta boxes in B, representing diffusing miRNAs in PBs and in the cytoplasm (Cyt) respectively. Scale bar, 1 μm. Dotted green circle represents PB outline in the first frame of the movie. Distribution of RNA (miRNA, mRNA and lncRNA diffusion constants in PB and Cyt are also depicted. Green area on the plot depicts the range of PB diffusion constants (n = 3, 55 cells). (H) Zoomed-in view of orange and violet boxes in C, from fixed UGD cells. Scale bar, 2 μm. Step-wise photobleaching trajectories PB- and Cyt-localized l7-Cy5/l7* is also shown. (I) Zoomed-in view of light yellow and red boxes in F, from fixed UGD cells. Scale bar, 2 μm. Single particle intensities of PB- and Cyt-localized FL-l7-6x-MS2 is also shown. (J) Distribution of RNA (miRNA, mRNA and lncRNA) abundance in PB and Cyt within fixed UGD cells (n = 3, 55 cells).

### RNAs stably or transiently localize at the core or periphery of PBs

To determine whether RNAs of distinct functionality interact stably or dynamically with PBs, we compiled the sub-granular dynamics and localization patterns of three main classes of RNAs (miRNAs, mRNAs and lncRNAs) and discovered hidden diversities in their PB interaction kinetics and localization patterns. By inspecting individual trajectories of RNAs localizing with PBs in live cells, we identified five distinct types RNA-PB interactions, each of which could be classified by a unique combination of diffusion coefficient (D), dwell time (T) and percentage of an RNA track colocalizing with a PB (P) (Figure 2A, S2A-B and Supplementary movies 2-6): 1) RNAs stably anchoring at PBs (D = 0.0001 – 0.1 μm^2^/s, T e 15 s, P = 100%, Supplementary movie 2); 2) RNAs displaying significant dynamics within PBs (D = 0.001 – 0.1 μm^2^/s, T ≥15 s, P = 100%, Supplementary movie 3); 3) RNAs entering PBs from the cytosol (D = 0.0001 – 0.01 μm^2^/s, T = 7.9 ± 0.7 s, P = 52 – 89%, Supplementary movie 4); 4) RNAs transiently probing PBs (D = 0.0001 – 1 μm^2^/s, T = 0.9 ± 0.1 s, P = 3 – 72%, Supplementary movie 5); and 5) RNAs exiting a PB into the cytosol (D = 0.0001 – 1 μm^2^/s, T = 0.8 ± 0.1 s, P = 7 – 83%, Supplementary movie 6). The first three and latter two interaction types depict what we refer to as stable and transient RNA-PB localizations, respectively. Using ratiometric single-molecule counting of RNAs in PBs and the adjacent cytosol (Figure S2C), and spatial mapping of RNPs with reference to PB boundaries (Figure S2D), we additionally discovered that RNAs were either clustered (enriched within PBs compared to the adjacent cytosol) or dispersed at PBs, and were found either localized near their core or their periphery/shell (Figure 2B-C) in fixed cells. Notably, the relative likelihoods of finding transient versus stable associations, clustered versus dispersed distributions, and PB-shell versus PB-core localizations was almost equal (Figure 2D). These findings identify diverse, dynamic modes of RNA-PB interactions, unraveling a new layer of complexity in the spatiotemporal organization of single RNA molecules during cellular homeostasis.

**Figure 2.**
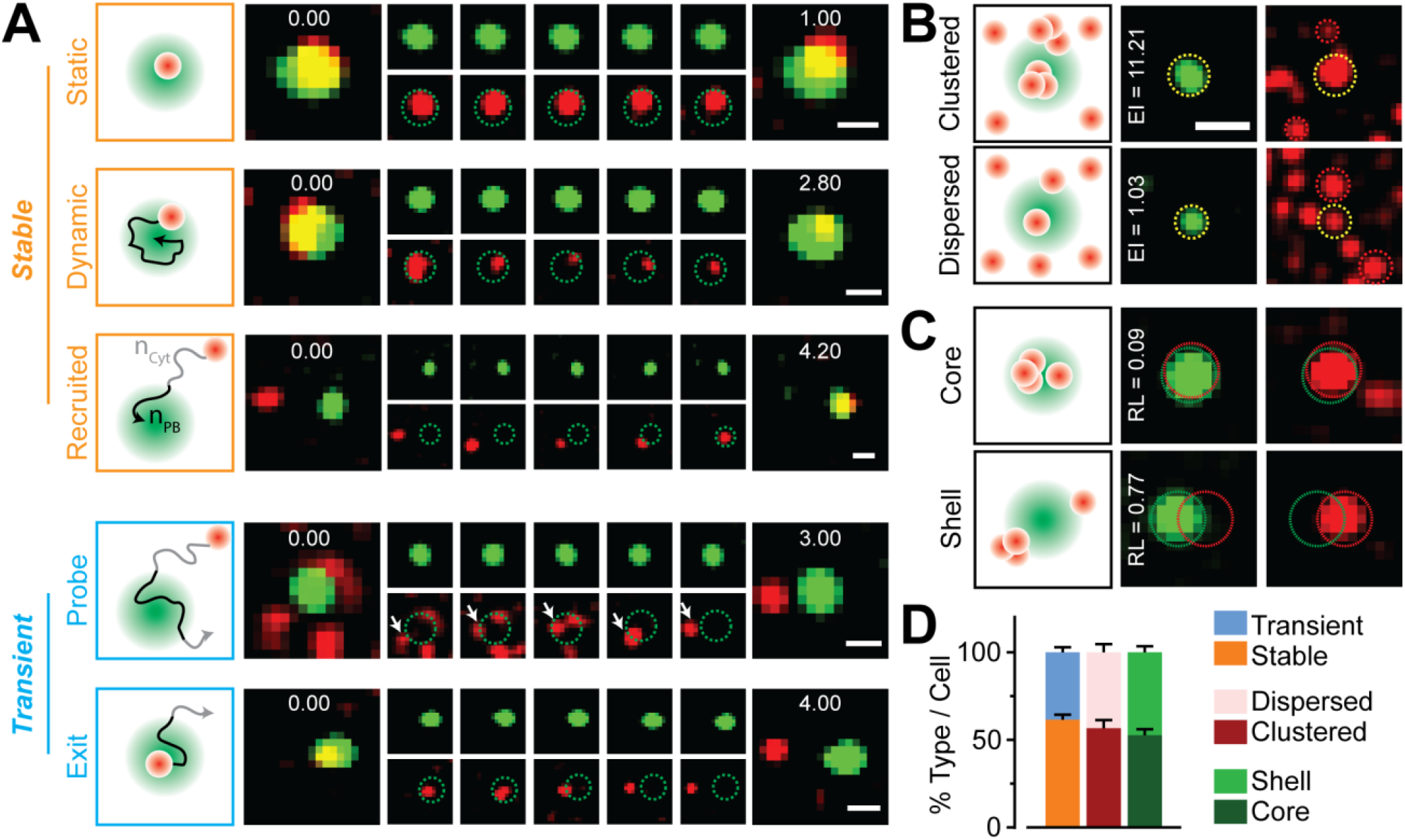
RNAs show diverse spatiotemporal localization patterns at PB core and periphery. (A) Schematic and representative time-lapsed images of PBs (green) and RNAs (red) in live UGD cells. Scale bar, 1 μm. Embedded numbers in green/red overlay images (far-left and far right) represent time in seconds. Dotted green circles in red panels have been included to aid in the identification of PB boundaries. White arrow points to an individual RNA particles. Stable RNA-PB association patterns (static, dynamic and recruited) are represented in orange whereas transient ones (probe and escape) are represented in blue. n_PB_ = number of track localizations within PBs, n_Cyt_ = number of track localizations in the cytosol. These two factors are used to calculate the track-wise percentage localization within PBs. (B) Schematic and representative images of PBs (green) and FL-l7-6x-MS2 mRNAs (red) representing the enrichment of RNAs in PBs within fixed UGD cells. Scale bar, 2 μm. Dotted yellow and red circles represent PB-RNA colocalization and cytoplasmic RNAs respectively. Enrichment index (EI) for these representative colocalizations are embedded in the green panels. (C) Schematic and representative images of PBs (green) and FL-l7-6x-MS2 mRNAs (red) representing the localization of RNAs within shells or cores of PBs in fixed UGD cells. Images are scaled as in B. Dotted green and red circles represent boundaries of PBs and RNAs respectively. Relative localization (RL) values of FL-l7-6x-MS2 mRNAs for these representative colocalizations are embedded in the green panels. (D) Percentage of stable vs transient, clustered vs dispersed, core vs shell RNA-PB colocalizations (n = 3, 110 cells).

#### lncRNA-PB interaction kinetics and patterns are distinct from those of miRNAs and mRNAs

To determine whether the general functionality of an RNA influences its interaction and localization with PBs, we compared in detail both live and fixed cell imaging data of l7-Cy5/l7* miRNA, FL-l7-6x-MS2 mRNA and THOR-MS2 lncRNA. While the fractional extent of RNA-PB colocalization did not significantly differ between the three classes of RNAs (Figure S3A), we found significant differences in the interaction kinetics and localization patterns between mi/mRNAs on the one hand and lncRNAs on the other (Figure 3). Firstly, THOR-MS2 displayed ~2-3-fold more transient PB interactions than l7-Cy5/l7* and FL-l7-6x-MS2 mRNA (Figure 3A). Although the photobleaching-corrected dwell time distributions were bi-phasic for all RNAs (Figure S3C), l7-Cy5/l7* (T_fast_ = 0.6 s and T_slow_ ≥ 15 s) and FL-l7-6x-MS2 (T_fast_ = 0.9 s and T_slow_ ≥ 15 s) resided at PBs for a significantly longer time than THOR-MS2 (T_fast_ = 0.6 s and T_slow_ = 2.9 s, Figure 3B and S3B). Additionally, we identified a third population of FL-l7-6x-MS2 and THOR-MS2 that dwelled at PBs for the entire duration of our observation window (15 s), without photobleaching, and hence were not captured by our dwell time calculations (Figure 3C). Notably, FL-l7-6x-MS2 displayed ~3-fold the relative fraction of such particles compared to THOR-MS2. We additionally found that THOR-MS2 frequently localized more to the shell of PBs than l7-Cy5/l7* and FL-l7-6x-MS2, which were ~2.5-5-fold more PB-enriched than THOR-MS2 and predominantly localized near PB cores (Figure 3D-E). Together, our data demonstrate that regulatory miRNAs and miRNA-regulated mRNAs infrequently escape PBs; by contrast, the lncRNA THOR only transiently associates with PBs, possibly due to IGF2BP1-mediated transcript stabilization (Hafner et al., 2010) and consistent with the relative dearth of lncRNAs in the transcriptome of PB cores (Hubstenberger et al., 2017).

**Figure 3.**
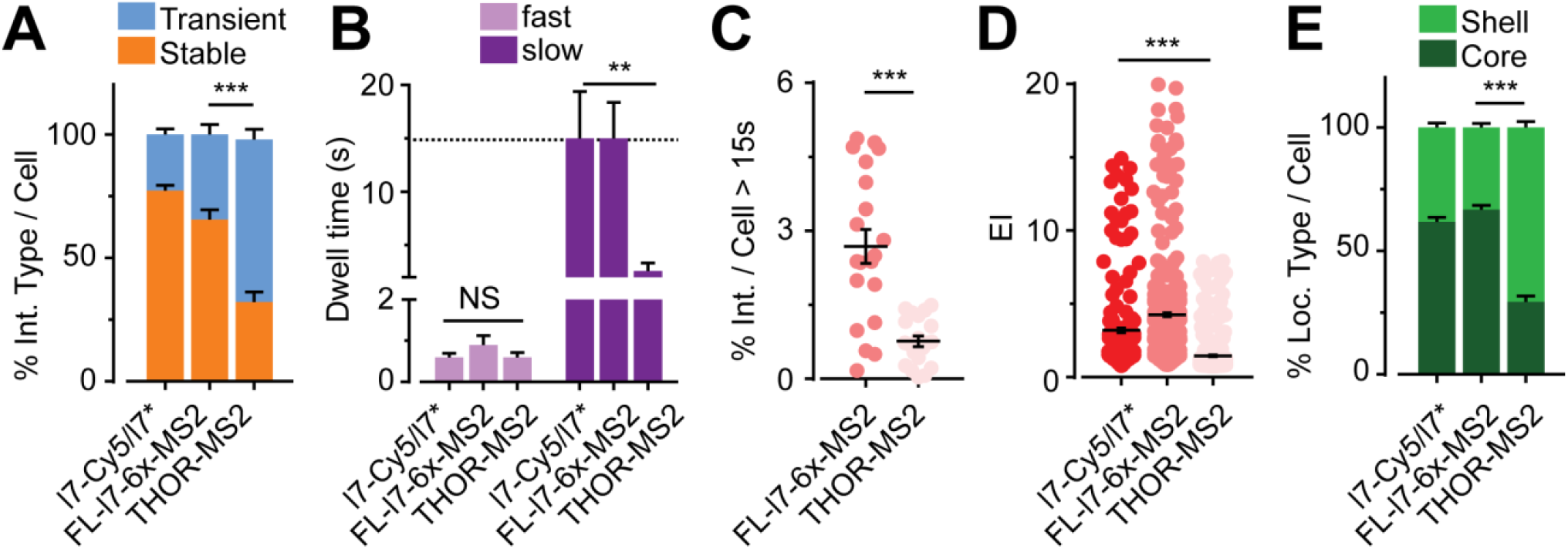
PB-interaction kinetics and patterns are different for distinct classes of RNAs. (A) Relative distribution of stable and transient interactions per live UGD cell for l7-Cy5/l7* miRNA, FL-l7-6x-MS2 mRNA and THOR-MS2 lncRNA. (B) Comparison of fast and slow RNA-PB interaction kinetics in live UGD cells. Dotted black line represents duration of acquisition. Photobleaching corrected dwell times that were greater than acquisition window were rounded to the acquisition time span. (C) Scatter plot representing % of RNA-PB interactions that last for the entire duration of imaging (15 s), without photobleaching per live UGD cell. (D) Scatter plot of EI per RNA-PB colocalization for different RNAs. Each dot represents an individual RNA-PB colocalization event in fixed UGD cells. (E) Relative distribution of core and shell localization patterns of different RNAs in fixed UGD cells. n = 3, 55 live and fixed cells respectively, NS, not significant, **p ≤ 0.001 or ****p ≤ 0.001 by two-tailed, unpaired Student’s t-test.

#### miRNA functionality influences miRNA-PB interaction kinetics and patterns

Next, we probed whether the specific RNA sequence within an RNA class influences its RNA-PB interactions. To this end, we compared the PB-localization of double-stranded l7-Cy5/l7* miRNA, whose regulatory functionality is clearly defined (Figure S1A), to that of two small RNA constructs and a small DNA construct of distinct intracellular stability (Pitchiaya et al., 2017) and regulatory potential: 1) l7/l7*-Cy5, i.e., let-7 miRNA Cy5-labeled on the passenger instead of the guide strand, where the passenger strand has very few endogenous targets and is at least 8-fold less stable than the guide strand; 2) ml7-Cy5/ml7*, a seed-sequence mutated let-7 miRNA variant that cannot bind endogenous let-7 targets and is at least 4-fold less stable than let-7 miRNA; and 3) dl7-Cy5/dl7*, a control DNA oligonucleotide of the same sequence as let-7 miRNA. As expected, the control DNA neither localized to, nor was enriched at PBs (Figure 4A-B). By contrast, the fractional extents of PB localization and enrichment were significant and similar for all three miRNAs (Figure 4A-B), suggesting that miRNA functionality is not necessary for PB localization. However, ml7-Cy5/ml7* and l7/l7*-Cy5 rarely displayed any transient interactions (Figure 4C), but instead exhibited monophasic dwell time distributions, residing in PBs for ≥ 15 s (Figure 4D and S4A). These observations suggest that transient PB interactions of a miRNA are correlated with its ability to repress mRNA targets, whereas unused (target-less) miRNAs are more stably recruited to PBs. Corroborating this notion, we found that, upon co-microinjecting its cognate (RL-ml7-2x) mRNA, the targeting ml7-Cy5/ml7* exhibited a substantial 5-fold increase in the fraction of transient interactions, resulting in a biphasic dwell time distribution with T_fast_ = 0.7 s and T_slow_ = 13.2 s (Figure 4E-F and S4C-D). Taken together, our results are consistent with PBs playing a role in surveillance for target-less miRNAs.

**Figure 4.**
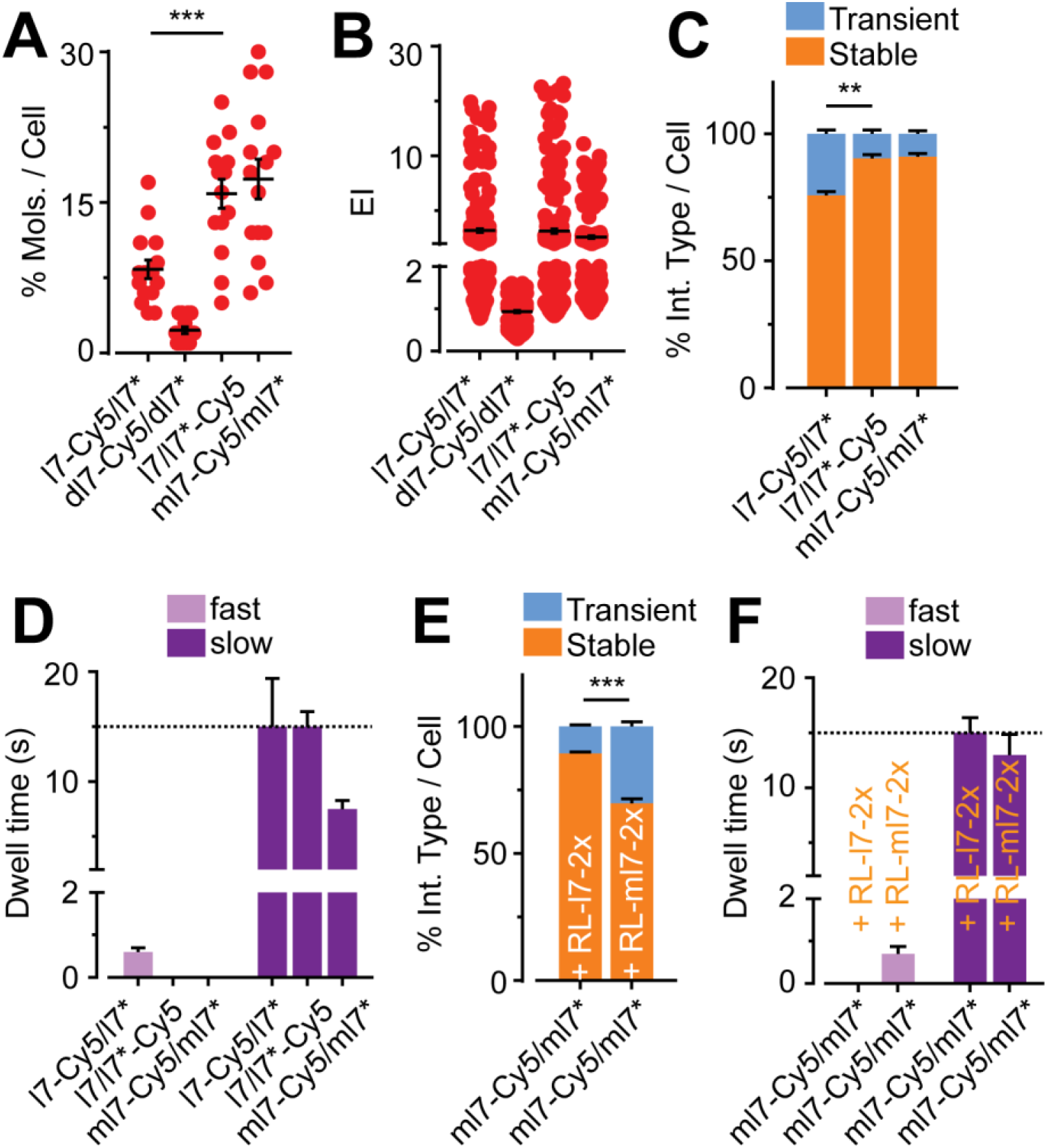
miRNA functionality influences miRNA-PB interaction kinetics and patterns. (A) Scatter plot representing the % of RNA or DNA molecules that colocalize with PBs per fixed UGD cell. Each dot represents a cell. (B) Scatter plot of EI for different constructs. Each dot represents an individual RNA/DNA-PB colocalization event in fixed UGD cells. (C) Relative distribution of stable and transient interactions per live UGD cell for different miRNAs. (D) Comparison of fast and slow miRNA-PB interaction kinetics in live UGD cells. (E) Relative distribution of stable and transient interactions per live UGD cell for ml7-Cy5/ml7* RNAs co-injected with a seed mismatched (RL-l7-2x) or seed matched (RL-ml7-2x) mRNA target. (F) Comparison of fast and slow ml7-Cy5/ml7*-PB interaction kinetics in the presence of a seed mismatched (RL-l7-2x) or seed matched (RL-ml7-2x) mRNA target in live UGD cells. n = 3, 15 cells per sample, **p ≤ 0.001 or ***p ≤ 0.0001 by two-tailed, unpaired Student’s t-test.

#### mRNA-PB interactions depend on translation potential and 3‘ versus 5‘terminal positioning of *cis*-regulatory elements

Given that we found miRNA recruitment to PBs to be dependent on their functionality, we next asked whether miRNA-regulated mRNAs behave similarly. To this end, we compared the PB localization dynamics of the let-7 regulated FL-l7-6x-MS2 mRNA (Figure S1) with those of FL-MS2 (lacking the regulatory 3‘UTR), FL-ml7-6x-MS2 (with a 3‘UTR containing six mutated let-7 MREs that are not targetable by endogenous let-7) and l7-6x-FL-MS2 (carrying a 5‘UTR with six tandem let-7 MREs, targetable by endogenous let-7) (Figure 5A). Notably, both FL-MS2 and FL-ml7-6x-MS2 were expressed to a much higher extent than the let-7-MRE containing FL-l7-6x-MS2 and l7-6x-FL-MS2 (Figure S5A). The latter two constructs were repressed to a similar extent by let-7 miRNA, suggesting that MREs embedded in either the 3‘ or 5‘UTR are both functional (Figure S5B). Strikingly, the fractional extents of localization and enrichment of l7-6x-FL-MS2 at PBs were similar to those of FL-MS2 and FL-ml7-6x-MS2, but significantly (at least 5-fold) lower than those of FL-l7-6x-MS2 (Figure 5B-C). Still, l7-6x-FL-MS2, FL-MS2 and FL-ml7-6x-MS2, much like FL-l7-6x-MS2, interacted transiently with PBs and displayed biphasic interaction kinetics (Figure 5D and S5C). While the "fast” phase for FL-MS2, FL-ml7-6x-MS2 and l7-6x-FL-MS2 (spanning ~0.5, 0.7 and 0.6 s, respectively), was similar to that of FL-l7-6x-MS2 (0.9 s), the "slow” phase for these constructs was ~3-fold shorter than that of FL-l7-6x-MS2 (3.2, 3.7 and 4.2 s, respectively, compared to 15 s, Figure 5E). A minority of FL-MS2, FL-ml7-6x-MS2 and l7-6x-FL-MS2 particles resided in PBs for the entire duration of acquisition (~ 15 s), but the number of such occurrences was ~4.5-fold lower than FL-l7-6x-MS2 (Figure 5F). These observations demonstrate that MRE inclusion in either the 3‘ or 5‘UTR induces mRNA repression; however, repression does not appear to correlate with PB localization. More specifically, our data imply that both unrepressed and 5‘UTR-repressed mRNAs dominantly exhibit transient PB interactions, whereas mRNAs with repressive MREs in the 3‘UTR have a higher propensity to bind factors that recruit them to PBs more stably.

**Figure 5.**
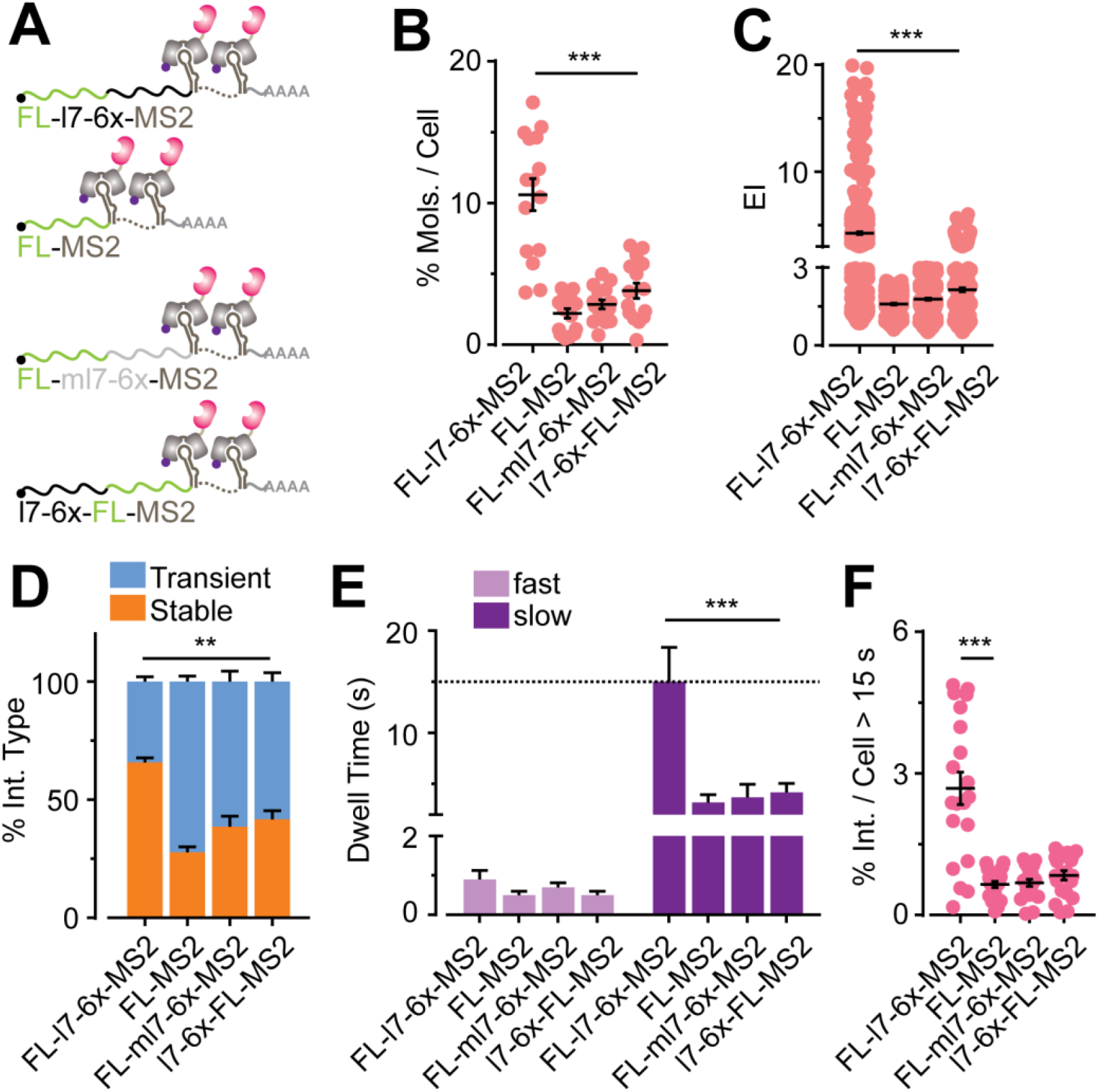
mRNA-PB interactions depend on translation potential and the 3‘ versus 5‘ position of MREs. (A) Schematic of four different mRNA constructs. Green line – CDS, black line – l7-6x MRE targetable by endogenous let-7, gray line – seed mutated ml7-6x MRE that cannot be targeted by endogenous let-7. MS2 stem loops and NLS-MCP-Halo as in Figure 1. (B) Scatter plot representing the % of mRNA molecules that colocalize with PBs per fixed UGD cell. (C) Scatter plot of EI for different mRNA constructs. Each dot represents an individual mRNA-PB colocalization event in fixed UGD cells. (D) Relative distribution of stable and transient interactions per live UGD cell for different mRNAs. (E) Comparison of fast and slow mRNA-PB interaction kinetics in live UGD cells. (F) Scatter plot representing % of RNA-PB interactions that last for the entire duration of imaging (15 s), without photobleaching per live UGD cell. n = 3, 20 cells per sample, **p ≤ 0.001 or ****p ≤ 0.001 by two-tailed, unpaired Student’s t-test.

#### mRNA turnover predominantly occurs outside of PBs

While a vast majority of our fluorophore labeled RNAs reside outside of microscopically visible PBs, seemingly inaccessible by the RNA processing enzymes enriched in these granules, a significant minority fraction localizes transiently or stably to PBs (Figure 2-5). Conversely, almost all visible PBs colocalize with RNA molecules, irrespective of relative RNA enrichment (Figure 6A), and a single PB encounters at least 3 RNA molecules under our imaging conditions (Figure 6B, S6B). Considering that RNAs and PBs frequently encounter each other and that PBs are enriched for RNA degradation enzymes (Hubstenberger et al., 2017; Parker and Sheth, 2007), we sought to test whether PBs are designated sites of RNA decay responsible for the bulk of cellular RNA turnover. While fluorescence microscopy can visualize large PBs (> 50 nm), it does not capture smaller functional complexes of RNA decay enzymes. We therefore kinetically modeled (Mourao et al., 2014) the mRNA degradation activity of microscopically visible and invisible PBs computationally (Figure 6C). We specifically tested miRNA-mediated mRNA decay, largely due its cellular prevalence and prior reports on miRNA programmed mRNA localization to PBs; however, our method is extendable to other decay processes. To this end, a two-dimensional lattice was populated with three molecular species, namely the effector miRNA induced silencing complex (miRISC), mRNP and PBs, whose abundances were based on prior reports (Aizer et al., 2014; Milo et al., 2010; Pitchiaya et al., 2012). We devised a basic set of reactions, each with predefined rates, whereby the interaction of miRISC with mRNPs activates PB-mediated mRNP degradation. Upon computing the copy number of each of these molecular species as they diffuse across the lattice through time, we found that mRNA degradation was most efficient when the lattice contained a large number of small, mobile PBs (Figure 6C). That is, while degradation is possible within large, microscopically visible PBs, the process is most efficient when degradation factors, perhaps individual molecules, are unconstrained in the cell, thus presenting a large surface area for capturing repressed mRNAs.

**Figure 6.**
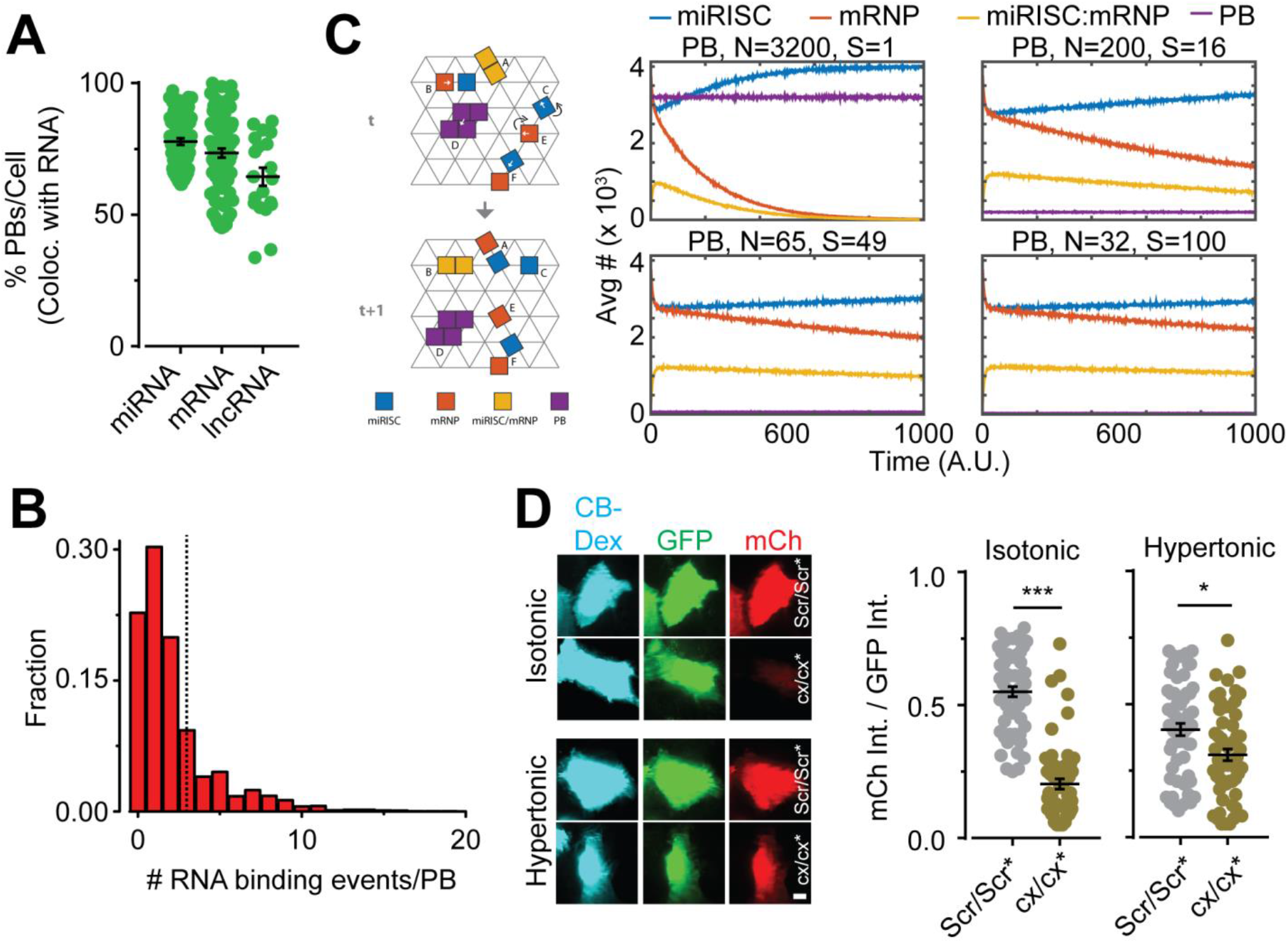
A majority of microscopically visible PBs associate with mRNAs, but mRNAs are more effectively degraded with a larger number of smaller, microscopically-invisible PBs. (A) Scatter plot representing the % PBs that colocalize with RNAs, per fixed UGD cell (n = 3, ≥ 15 cells per sample). (B) Frequency distribution of the number of times an individual PB encounters an RNA in live UGD cells (n = 3, 175 cells, 2742 PBs). Dotted line represents the average number of RNA encounters per PB after correcting for photobleaching. (C) Schematic (left) of *in silico* kinetic modeling of RNA-PB interactions and RNA decay. Changes in the abundance of each molecular species (represented in schematic) over the timescale of the simulation is also depicted (right). About a quarter of miRISC molecules initially bind to mRNP molecules to form miRISC:mRNP complexes. As mRNPs are depleted, the number of miRISC:mRNP complexes and miRISC gradually decrease and increase, respectively. With 3200 P-bodies of size 1 each (top left), all mRNP molecules are depleted during 1000 iterations of the simulation (red curve). With 200 P-bodies of size 16 each (top right), < 1500 mRNP molecules persist after 1000 iterations. With 65 P-bodies of size 49 each (bottom left), 2000 mRNP molecules persist after 1000 iterations. With 32 P-bodies of size 100 each (bottom right), akin to the abundance and size of microscopically visible PBs, ~ 2250 mRNP molecules persist after 1000 iterations. (D) Experimental validation of simulations using microinjection-based miRNA activity assay. Left, representative images of U2-OS cells treated with isotonic or hypertonic (300 mM Na+) medium and co-injected with CB-Dextran, GFP mRNA,mCh mRNA with MREs for cxcr4 (cx/cx*) miRNA and either a scrambled, control siRNA (Scr/Scr*) or cx/cx*. Images were acquired 4 h after injection. Right, scatter plot representing the ratio of mCh: GFP intensity at various injection and treatment conditions. Each dot represents a U2-OS cell (n = 3, 60 cells for each sample).

To test our *in silico* predictions intracellularly, we resorted to modulating PB number and size via hyperosmotic stress, a method that has been proven to increase PB number in yeast (Huch and Nissan, 2017). We confirmed that hyperosmotic treatment of UGD cells results in a high number of immobile GFP-Dcp1a foci (Figure S6B-D). Microinjection-based miRNA activity assays (Figure S1E) in U2-OS cells suggested that, as predicted, miRNA-mediated gene repression is alleviated when PBs are aggregated upon subjection of cells to hyperosmotic stress (Figure 6D). Taken together, our data support the notion that mRNA degradation is primarily mediated by degradation enzymes freely diffusing in the cytosol, relegating PBs to degrading only a small fraction of repressed mRNAs.

### DISCUSSION

Previous reports have provided exquisite static snapshots of RNA and protein colocalization with PBs (Cougot et al., 2012; Horvathova et al., 2017; Kedersha and Anderson, 2007; Liu et al., 2005), but could not assess the dynamics of the underlying recruitment processes. Others have provided valuable information regarding the dynamics of PB movement and the bulk exchange of proteins or mRNAs between PBs and the cytosol, but could not extract mechanistic information about the recruitment of biomolecules to PBs (Aizer et al., 2008; Aizer et al., 2014; Kedersha et al., 2008; Leung et al., 2006). Using single-molecule live-cell imaging we here uniquely demonstrate that miRNAs, mRNAs and lncRNAs dynamically localize to PB either stably or transiently (Figures 1 and 2). Stable anchoring at PBs is concordant with prior snapshots that visually portray RNA accumulation within PB “cores”, whereas more mobile localizations and transient interactions are more likely to depict the localization of RNAs in PB “shells” (Cougot et al., 2012). In agreement with our data on mi/m/lncRNA-PB interactions during cellular homeostasis, recent reports (Moon et al., 2018; Wilbertz et al., 2018) have complementarily shown that mRNAs associate both stably and transiently with both stress granules (SGs) and PBs during the integrated stress response. The dwell times annotated as stable (~250 s) or transient (~10 s) in these reports are akin to particles in our datasets that dwell at PBs for the entire duration of acquisition (> 15 s) and for ~3-5 s, respectively. We have found an additional, highly dynamic interaction mode that lasts < 1 s, which potentially represents a relatively rapid PB-probing step. Upon RNP remodeling, these rapid encounters may transition into longer spans of granule probing or stable docking of RNAs to granules.

Elucidation of the PB-core transcriptome (Hubstenberger et al., 2017) has suggested that certain miRNAs, lncRNAs and repressed mRNAs are enriched in PBs, yet it is unclear whether the principles governing PB enrichment for these major classes of transcripts are similar or different. Strikingly, we found that miRNAs, mRNAs and lncRNAs have distinct PB localization signatures, which appear correlated with the distinct functionalities of these transcripts and the diversity in the types of RNPs they form (Figure 3). Based on our data, we propose a model that assigns PB localization patterns to specific RNA forms and functionalities (Figure 7). Stably anchored and PB-enriched miRNAs are predominantly dysfunctional – they do not have many mRNA targets and localize to PBs in their unbound or miRISC-bound (single-stranded or double-stranded) forms (Figure 4). Functional miRNAs, more likely to be in RISC-mRNA complexes, display this behavior only in their minority and, when anchored, preferably localize within PB cores. These data are consistent with prior reports that both strands of both target-less and target-containing siRNAs localize to PBs (Jakymiw et al., 2005). We posit that, by contrast, transient associations at PB peripheries represent miRISC-mRNA complexes that do not yet have bound an important recruiting protein, such as GW182 or LAMP1 (Moon et al., 2018; Wilbertz et al., 2018), that is required for PB association. Conversely, highly translatable mRNAs that are not associated with miRNAs, while transiently associating peripherally, are not enriched at PBs. Based on recent reports (Moon et al., 2018; Wilbertz et al., 2018) and our data (Figure 5) we predict that non-translating mRNAs and translationally repressed mRNAs bearing MREs in their 3‘UTR stably associate with PB cores, while only the latter are enriched at PBs. Furthermore, we find that miRNA-repressed mRNAs with MREs in the 5‘UTR (Figure S5A-B) are not enriched at and only transiently associate with PBs, probably also due to the lack of a PB-recruitment factor bound to these RNPs (Figure 5). Prior reports have demonstrated that MREs in the 5’ UTR cause translational repression downstream of translation initiation sites (Lytle et al., 2007), potentially resulting in polysome bound non-translating mRNAs, which consequently cannot enter ribosome excluded PB cores (Parker and Sheth, 2007). By contrast, MREs in the 3’UTR typically result in inhibition of translation initiation, leading to non-translating mRNAs that are also free of ribosomes, which can then enter PB cores. Taken together, our data suggest that different modes of miRNA-mediated mRNA repression favor different types of PB localization. Our data also suggest that polysome association negatively impacts stable PB localization, consistent with observations that actively translating mRNAs only transiently associate with SGs (Moon et al., 2018; Wilbertz et al., 2018).

**Figure 7.**
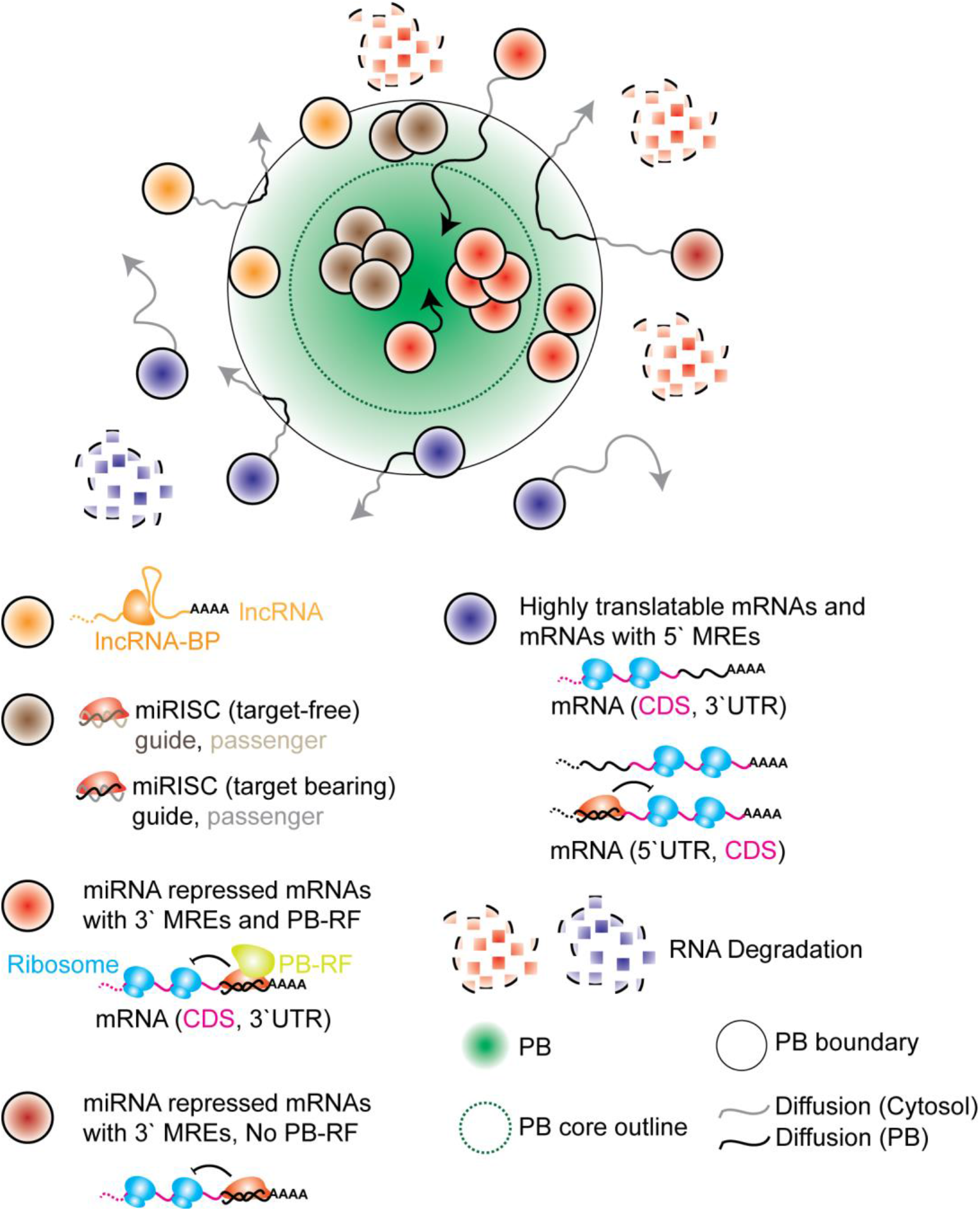
Resulting model for the dynamic recruitment of specific RNAs to PBs. RNAs dynamically associate with PB core or shell based on functionality. Target-free miRNAs, mRNA-targeting miRNAs and miRNA-targeted mRNAs with 3‘UTR MREs are stably enriched within either cores or shells of PBs. The presence of a PB recruitment factor (PB-RF) may influence the dynamics and enrichment extent of miRNA-targeted mRNAs at PBs. lncRNAs, translating mRNAs and miRNA-targeted mRNAs with 5‘UTR MREs transiently associate with PB shells. A majority of nuclease mediated RNA degradation occurs outside of PBs.

THOR is a recently discovered, highly conserved testis-specific lncRNA that is up-regulated in a broad range of human cancers and found to work in concert with IGF2BP1 (Hosono et al., 2017), a PB-enriched protein (Hubstenberger et al., 2017), to potentially stabilize various transcripts via coding region instability determinant-mediated mRNA stabilization (Weidensdorfer et al., 2009). Based on our data, we propose that functional THOR assembles into slowly diffusing (D = 0.0001 – 0.1 μm^2^/s) RNPs that mediate its function (Hosono et al., 2017). The frequent, transient associations with PBs may be linked to the regulatory role of this lncRNA, wherein one can envision: 1) THOR depositing regulated mRNAs for storage at PBs; or 2) THOR instead selecting PB-stored mRNAs for reintroduction into the translating cytoplasmic pool. Of note, we rarely observed any stable anchoring or enrichment of THOR at PBs, which suggests that the mere inability of RNA to be translated is not a sole prerequisite for stable PB association and enrichment.

Our molecular observations of colocalizations can be explained by phase transition principles that recently have been recognized to govern the assembly of large membrane-free granules (Protter and Parker, 2016; Shin and Brangwynne, 2017). Static, core-localized and enriched RNPs may serve as nucleating factors for large PBs, whereas dynamic, shell-localized and dispersed colocalizations may occur when the interfaces of the RNP, PB and surrounding cytoplasm are similar, as in a Neumann's triangle observed in Cajal bodies attached to B-snurposomes (Shin and Brangwynne, 2017). Transient colocalizations may represent cases where the smaller RNP and PB come in close proximity, but the interfacial surface tension is too high for the two to fuse, presumably due to the absence of an appropriate PB-recruitment factor on the RNP.

Although there is a general agreement on the phase-separation assembly principles of PBs and other RNA granules, the functions of these granules are a topic of intense debate. Certain reports have suggested that PBs may have stress dependent RNA decay or storage roles (Aizer et al., 2014), whereas others have suggested that PBs are sites of RNA storage, but not decay (Eulalio et al., 2007b; Horvathova et al., 2017; Stalder and Muhlemann, 2009; Tutucci et al., 2017). Notably, all previous studies have examined only microscopically visible PBs. Our computational simulations, which considered PBs both large and small, together with subsequent experiments using hyperosmotic stress to induce PB aggregation, suggest that microscopically visible PBs cannot account for the bulk of cellular mRNA decay (Figure 6). Our data instead suggest that fundamental principles of physical chemistry hold true for mRNA decay processes within the complex cellular environment, in that the entropic gain from the larger degree of freedom and surface area of freely diffusing decay components dominates. In addition to storing repressed mRNAs, our work unveils an additional housekeeping role for PBs in storing or possibly degrading unused miRNAs for their surveillance. Super-resolved fluorescence microscopy thus is shown to provide a powerful approach for mechanistically probing the dynamic assembly of RNP granules by phase separation at single-molecule resolution.

## ACKNOWLEDGEMENTS

We thank T.C. Custer and M. Denies for technical assistance. We thank N. Kedersha, D. Bartel and R. Singer for generous gifts of plasmids containing GFP-Dcp1a, the 3‘UTR of HMGA2 and the MS2 system of plasmids respectively. We thank L. Lavis for providing JF646 dye. This work was supported by National Institutes of Health (NIH) R01 grant GM081025 and a University of Michigan Comprehensive Cancer Center/Biointerfaces Institute Research Grant to N.G.W., whereas A.J. is supported by the NIH Cellular and Molecular Biology Training Grant T32-GM007315. A.M.C. is supported by the Prostate Cancer Foundation, by the Howard Hughes Medical Institute and is an American Cancer Society Research Professor. S.P. was supported by an AACR-Bayer Prostate Cancer Research Fellowship (16-40-44-PITC). L.X. was supported by a US Department of Defense Postdoctoral Fellowship (W81XWH-16-1-0195). We also acknowledge NSF MRI-ID grant DBI-0959823 to N.G.W. for seeding the Single Molecule Analysis in Real-Time (SMART) Center whose Single Particle Tracker TIRFM equipment was used for much of this study.

## AUTHOR CONTRIBUTION

S.P. designed and performed all assays. M.D.A.M. and S.S. performed the kinetic modeling. A.J. assisted with immunofluorescence assays. L.X. and X.J. created and validated mRNA and lncRNA constructs. S.P., S.S., A.M.C. and N.G.W. conceived the study and all authors wrote the manuscript together.

## METHODS

### DNA, RNA and LNA oligonucleotides

All DNA and RNA oligonucleotides used for iSHiRLoC experiments and reverse transcription, followed by quantitative polymerase chain reaction (RT-qPCR) were obtained from IDT. Oligonucleotides contained a 5‘ Phosphate (P) and, in the case of fluorophore labeled oligonucleotides, a Cy5 dye at the 3‘end. Dyes were attached after oligonucleotide synthesis to a 3‘amino group on a C6 carbon linker and were HPLC purified by the vendor. Guide and passenger strands were heat-annealed in a 1:1.1 ratio to achieve 10 μM stock solutions and were frozen until further use. Negative control siRNA (Scr/Scr*) was purchased as ready-to-use duplex samples from Ambion respectively. Six tandem let-7 (l7-6x) miRNA response elements (MREs) or mutant l7-6x (ml7-6x) MREs were purchased as gene blocks from IDT. AntimiR LNA oligos were purchased from Exiqon. Oligonucleotide and gene block sequences are listed in Table S1.

### Plasmids

pEGFP-Dcp1a was constructed by ligating PCR amplified (using Pfu ultra polymerase, Agilent, # 600380) EGFP ORF (from pEGFP-C1, Clontech) into pmRFP1-hDcp1a (gift from Nancy Kedersha, Brigham Women‘s hospital) within the AgeI and XhoI restriction enzyme (RE) sites. This replaces mRFP1 with EGFP in the plasmid. pEF6-mCh and pEF6-mCh-cx-6x construction was previously described previously (Pitchiaya et al., 2017). pEF6-mCh-l7-6x plasmid was constructed by ligating l7-6x gene block within NotI and XbaI sites of pEF6-mCh plasmid. Plasmids pRL-TK-let7-A, pRL-TK-let7-B, pRL-TK-cxcr4-6x, phage-ubc-nls-ha-2xmcp-HALO (a gift from Phil Sharp, Addgene plasmid # 11324, #11325, # 11308 and # 64540) and pmiR-GLO (pmG, Promega, # E1330) were purchased. pmG-MS2, encoding the firefly luciferase (FL) gene followed by 24 MS2 stem loops (FL-MS2), was created in two steps. First, the coding sequence (CDS) of IF2 was PCR amplified and ligated into the SbfI and NotI RE site of pmG, to create pmG-IF2. MS2 stem loops from pSL-MS2_24x (a gift from Robert Singer, Addgene plasmid # 31865) were then cloned into the EcoRI (introduced by above PCR)-NotI restriction enzyme sites pmG-IF2, to generate pmG-MS2. Clones containing the MS2 stem loops were created in SURE2 bacterial cells (Stratagene) to minimize recombination of the MS2 repeats with the bacterial genome. pmG-l7-6x-MS2 and pmG-ml7-6x-MS2 encoding FL-l7-6x-MS2 and FL-ml7-6x-MS2 respectively were constructed by ligating the l7-6x or ml7-6x gene blocks within the XhoI RE site in pmG-MS2. l7-6x-pmG-MS2 encoding l7-6x-FL-MS2 was created using Gibson ligation of the l7-6x gene block between the human phosphoglycerate kinase promoter and FL CDS. pLenti6-THOR and pLenti6-RHOT (antisense of THOR) were constructed as described (Hosono et al., 2017). plenti6-THOR-MS2 was constructed by ligating 24x MS2 stem loops from pmG-MS2 into EcoRI and NotI sites of pLenti6-THOR.

### Cell culture

U2OS (HTB-96, ATCC) cells were propagated in McCoy’s 5A (GIBCO, # 16600) supplemented with 10% FBS (GIBCO, # 16000). U2-OS cells stably expressing GFF-Dcp1a (UGD) was created by transfecting U2-OS cells with pEGFP-Dcp1a and selecting for stable clones by G418 selection. UGD cells were grown in the abovementioned medium supplemented with 100 μg/mL G418 (Thermo-Fisher, # 10131027). All medium typically contained 1x Penicillin-Streptomycin (GIBCO, #15140). Phenol-red free McCoy’s 5A (GE-Amersham, # SH3027001) supplemented with 1% FBS was used for seeding and cells for imaging experiments. For hyperosmotic shock, cells were treated with the above media supplemented with 10 x PBS such that the final sodium concentration was 300 mM. Plasmid transfections for MS2-MCP imaging and cell growth assays were achieved using Fugene HD (Promega, # E2311). Cotransfection of plasmids with oligos was achieved using lipofectamine 2000 (ThermoFisher, #11668027)

### mRNA synthesis

pRL-TK-cx6x, pRL-TK-let7-A and pRL-TK-let7-B were linearized with NotI to generate RL-cx6x RL-l7-2x and RL-ml7-2x mRNAs respectively. pEF6-mCh-cx6x and pEF6-mCh-l7-6x were linearized with XbaI to generate mCh-cx6x and mCh-l7-6x mRNA respectively. The pCFE-GFP plasmid (Thermo Scientific) was directly used in the in vitro transcription reactions to generate the GFP mRNA. The linearized plasmids were extracted with phenol and chloroform and subsequently ethanol precipitated. In vitro transcriptions were performed using the MegaScript T7 kit (Thermo-Fisher, # AM1334) according to manufacturers protocol. Transcription reactions were then DNase treated (turbo DNase supplied with kit) and the respective RNAs were purified by sequential gel-filtration chromatography (Nap-5 followed by Nap-10, GE healthcare, # 17085301 and #17085401 respectively) and ethanol precipitation. The RNAs were 5’capped (ScriptCap™ m7G Capping System, CELLSCRIPT, # C-SCCE0625) and polyadenylated (A-Plus™ Poly(A) Polymerase Tailing Kit, CELLSCRIPT, # C-PAP5104H) and were further purified by sequential gel-filtration chromatography and ethanol precipitation. The length of the polyA tails was estimated based on electrophoretic mobility on a 1.2% formaldehyde agarose gel.

### Luciferase reporter assays

100 μL of 10, 000 −20, 000 cells were seeded per well of a 96 well plate in antibiotics-free medium. Transfection conditions and luminescence readouts are as described previously (Pitchiaya et al., 2012; Pitchiaya et al., 2013; Pitchiaya et al., 2017). Briefly, cells were transfected with 60 ng of the indicated plasmid, 10 nM of the indicated dsRNA, and when appropriate 30 nM anti-ctrl or anti-l7 antimiRs, 0.4 μL of Lipofectamine 2000 (Invitrogen) and 50 μL of OptiMEM (GIBCO). 6 h after transfection the growth medium was replaced with fresh medium. 24 h after transfection, medium was replaced with phenol red-free McCoy’s 5A. Dual luciferase assays were performed using the Dual-Glo luciferase assay reagents (Promega, # E2920) as per the manufacturer’s protocol and luminescence was detected using a Genios Pro (Tecan) plate reader.

### RT-qPCR

Cells were harvested and total RNA from cells were isolated using QIAzol Lysis reagent (Qiagen) and the miRNeasy kit (Qiagen) with DNase digestion according to the manufacturer’s instructions. cDNA was synthesized using Superscript III (Invitrogen) and random primers (Invitrogen). Relative RNA levels determined by qRT-PCR were measured on an Applied Biosystems 7900HT Real-Time PCR System, using Power SYBR Green MasterMix (Applied Biosystems). Expression was quantified by 2^ΔCt^ method, wherein Myc expression was first normalized to that of GAPDH and then this normalized expression was further normalized to Mock treatment.

### Cell growth assays

100 μL of 10, 000 −20, 000 cells were seeded per well of a 96 well plate in antibiotics-free medium and were transfected every 24 h with the appropriate plasmid construct using Fugene HD (Promega, # E2311). Cell growth and viability was measured as an end point measurement for each time point using the Cell-titer GLO assay (Promega, # G7570) based on manufacturer’s instructions.

### Microinjection

Cells grown on DeltaT dishes (Bioptechs, # 0420042105C) were microinjected as described (Pitchiaya et al., 2012; Pitchiaya et al., 2013, Pitchiaya et al., 2017). Briefly, injection solutions contained the appropriate miRNA 1 μM concentration, 1x PBS and 0.5 mg/mL of 10 kDa cascade blue conjugated dextran (CB-Dex, ThermoFisher, # D1976). For microinjection based miRNA activity assay, mRNAs were added at a stoichiometric amount based on the number of miRNA binding sites, for instance, 0.16 μM of RL-cx6x mRNA, bearing 6 cxcr4 binding sites, was added along with 1 μM cxcr4 miRNA. Solutions were filtered through a 0.45 μm Ultrafree-MC filter (Millipore, # UFC30HV00) and then centrifuged at 16,000 x g for 15min at 4 °C immediately before injection. The solution was loaded into a femtotip (Eppendorf, # E5242952008). Injections were performed using a Femtojet pump (Eppendorf) and an Injectman (Eppendorf) mounted to the microscope. Microinjections were performed at 100 hPa injection pressure for 0.5 s with 20 hPa compensation pressure. This pressure translates to a volume of 0.02 pL and 10,000-20,000 miRNA molecules.

### Single-molecule fluorescence *in situ* hybridization

smFISH was performed as described (Hosono et al., 2017). Briefly, cells were grown on 8-well chambered coverglasses (Thermo-Fisher, # 155383PK), formaldehyde fixed and permeablized overnight at 4 ^°^C using 70% ethanol. Cells were rehydrated in a solution containing 10% formamide and 2 × SSC for 5 min and then treated with 100 nM fluorescence in situ hybridization probes (LGC-Biosearch) for 16 h in 2 × SSC containing 10% dextran sulfate, 2 mM vanadyl-ribonucleoside complex, 0.02% RNAse-free BSA, 1 μg μl-1 E. coli transfer RNA and 10% formamide at 37 °C. After hybridization, cells were washed twice for 30min at 37 °C using a wash buffer (10% formamide in 2 × SSC). Cells were then mounted in solution containing 10 mM Tris/HCl pH 7.5, 2 × SSC, 2 mM trolox, 50 μM protocatechiuc acid and 50 nM protocatechuate dehydrogenase. Mounts were overlaid with mineral oil and samples were imaged immediately. Sequences of Q670 labeled probes against the FL gene are listed in Table S1 and probes against THOR were previously described (Hosono et al., 2017).

### Immunofluorescence

Cells were grown on 8-well chambered coverglasses (Thermo-Fisher, # 155383PK), formaldehyde fixed and permeablized using 0.5% Triton-X100 (Sigma, T8787-100ML) in 1x PBS at room temperature (RT) for 10 min. Cells were then treated with blocking buffer containing 5% normal goat serum (Jackson Immunoresearch, 005-000-121), 0.1% Tween-20 (Sigma, P9416-50ML) in 1x PBS at RT for 1 h. Primary antibodies (pA) were diluted in blocking buffer to appropriate concentrations and cells were treated with pA at RT for 1 h. Following three washes with the blocking buffer for 5 min each cells were treated with secondary antibodies (sA) diluted in blocking buffer to appropriate concentrations. Following two washes with the blocking buffer and two washes with 1x PBS for 5 min each, cells were mounted in solution containing 10 mM Tris/HCl pH 7.5, 2 × SSC, 2 mM trolox, 50 μM protocatechiuc acid and 50 nM protocatechuate dehydrogenase. Mounts were overlaid with mineral oil and samples were imaged immediately. We used the following antibodies in our study: rabbit-anti-Dcp1a (Sigma, HPA013202-100UL, 1:100), rabbit-anti-Dcp2 (Thermo, PIPA5-34455, 1:200), rabbit-anti-Xrn1 (Bethyl labs, A300-443A, 1:200), rabbit-anti-DDX6 (Bethyl labs, A300-460A, 1:200), mouse-anti-GW182 (Abcam, ab70522, 1:150), anti-Ago2 (Sino biological, 50683-RO36, 1:200) and rabbit-anti-CNOT1 (Proteintech, 14276-1-AP, 1:50).

### Microscopy

Highly inclined laminated optical sheet (HILO) imaging was performed as described (Pitchiaya et al., 2012; Pitchiaya et al., 2013, Pitchiaya et al., 2017) using a cell-TIRF system based on an Olympus IX81 microscope equipped with a 60x 1.49 NA oil-immersion objective (Olympus), as well as 405 nm (Coherent ©, 100 mW at source, ~65 μW for imaging CB-Dex), 488 nm (Coherent ©, 100 mW at source, ~1.2 mW for imaging GFP), 561 nm (Coherent ©, 100 mW at source, ~50 μW for imaging mCh) and 640 nm (Coherent ©, 100 mW at source, 13.5 mW for imaging Cy5) solid-state lasers. Quad-band filter cubes consisting of z405/488/532/640rpc or z405/488/561/640rpc dichroic filters (Chroma) and z405/488/532/640m or z405/488/561/640m emission filters (Chroma) were used to filter fluorescence of the appropriate fluorophores from incident light. Emission from individual fluorophores was detected sequentially on an EMCCD camera (Andor IXon Ultra) for fixed cell imaging. For multicolour live-cell imaging, the emitted light was split onto two different EMCCDs using a single beamsplitter within a filter adapter (TuCam, Andor). Emission filters were placed just prior to each camera to minimize fluorescence bleed-through. For simultaneous detection of GFP and Cy5, a filter set with a 585dxcru dichroic that splits fluorescence into et525/50m and et705/100m emission filters respectively was placed in the Tucam adapter. For live cell imaging of MS2-MCP constructs, UGD cells on Delta T dishes were treated with 100 nM JF646-Halo ligand (a kind gift from Luke Lavis) for 30 min in growth medium without phenol red (Grimm et al., 2015). After the treatment, cells were washed three times in media and placed back in the incubator for 30 min, prior to imaging.

### Image Analysis

The two cameras used for simultaneous acquisition of GFP and Cy5 fluorescence in live cells were first registered as described (Churchman et al., 2005). Registration was achieved by imaging 0.1 μm tetraspeck beads (Thermo-Fisher, # T7279), whose emission is similar to both GFP and Cy5, before or after imaging of live cells. The registration matrix was then applied to GFP and Cy5 images for accurate tracking of PBs and RNAs respectively. Single particle tracking was performed as described (Pitchiaya et al., 2012; Pitchiaya et al., 2013) with some minor modifications. Briefly, particle tracking analysis was performed in Imaris (Bitplane) using tracks that spanned at least four video frames and all tracks were fit to a Brownian diffusion model to extract diffusion coefficients. PB boundaries were detected using a local contrast/threshold approach in Image J and Imaris. An RNA particle was identified as colocalizing with a PB when the centroid of the RNA is at or within the boundary of a PB. The use of finite observation windows to measure the dwell times introduces a systematic bias in the observed dwell times. This was corrected for by measuring the aggregate time for Cy5 photobleaching (T_phb_) and subtracting its reciprocal this from the reciprocal of the observed dwell time (T_obs_) along with the reciprocal of the observation window (T_w_), as described by T_actual_ = 1 / ((1/ T_obs_) - (1/ T_phb_) - (1/ T_w_)). Percentage of track colocalizing with PBs (track %) was calculated as n_PB_ / (n_PB_ + n_Cyt_), where n_PB_ = number of track localizations within PBs, n_Cyt_ = number of track localizations in the cytosol and depicted in Figure 2.

Step-wise photobleaching analysis of fluorophore labeled miRNAs and intensity analysis of smFISH particles in fixed cells were done using custom written Lab-view codes and ImageJ macros that can be shared upon request, as described (Pitchiaya et al., 2012; Hosono et al., 2017). To overcome statistical biases of co-incidental colocalizations introduced merely by RNA abundance, we calculated the accumulation of RNA within PBs via an enrichment index (EI) – a ratio of the number of RNA molecules in PB to those outside of PBs (Figure 2 and S2). An E.I. of > 1 suggests that the RNA accumulates at PB, whereas the opposite is true if the E.I. is ≤ 1. Relative localization (RL) of RNAs within PBs was calculated as d_CR_ / (d_RB_ + d_CB_), where d_CR_ = distance of RNA centroid from PB centroid, d_RB_ = distance of RNA centroid from PB boundary, d_CB_ = distance of PB centroid from PB boundary and depicted in Figure S2.

mCh and GFP signal from microinjection based miRNA activity assay were extracted and analyzed as described (Pitchiaya et al., 2012; Pitchiaya et al., 2017). Briefly, mCh and GFP intensity threshold were set (Huang threshold in image J) to automatically identify cell boundary. Background intensity, outside of cell boundary, was subtracted from mCh and GFP signal to extract the corrected intensity, whose ratio was calculated on a per cell basis.

### *In silico* kinetic modeling

The fundamental theory and basic methodology of modeling, including the lattice gas automata algorithm are as described (Mourao et al., 2014). Our simulation platform allows for the specification of a variable number of elementary reactions. Unless otherwise stated, the results presented here were obtained using two different reactions:

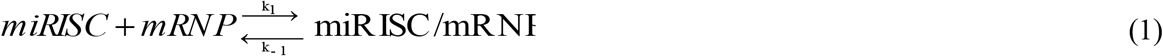

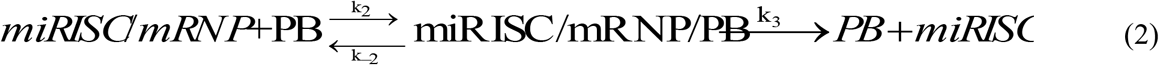

The reaction in (2) represents a catalytic event. The rate coefficients k_i_ are modeled as reaction probabilities. For example, in (1) k_1_ is modeled by the probability that a miRISC and an mRNP molecule will react to form complex miRISC/mRNP, given that they have collided. Unless otherwise stated, the probability of a forward reaction (on the basis of the rate coefficients k_1_ and k_2_) is set to 1 and the probability of a reverse reaction (on the basis of the rate coefficients k_-1_ and k_-2_) is set to 0.1. The probability of a catalytic reaction (on the basis of the rate coefficient k_3_) is set to 0.1. Note that the forward reaction rates (e.g., k_1_) may remain constant over time, in agreement with the law of mass action, or decay over time for diffusion-limited reactions, when the time required for any two reactants to interact increases with the level of obstruction to diffusion. In the latter case, it can be shown that log(k_1_) decays linearly at long times in a logarithmic time scale, as described (Mourao et al., 2014).

Each simulation begins with all particles randomly placed on a 2D lattice of size 200×200 lattice points with cyclic boundary conditions. Particles can be initialized with different sizes, provided that they are square, i.e., each initial particle can only occupy x2 positions, x being at least 1. Our platform allows for the creation of initial aggregates of a particular number and size. With the restriction mentioned above, we modulate the number and size of P-body particles within an aggregate with the assumption that all P-body particles within an aggregate have the same size. Each aggregate of P-bodies is created in two main steps. In the first step occurs, we insert the first molecule of the aggregate in the lattice. This first molecule is placed in a random position in the lattice. In the second step, we randomly select an adjacent neighborhood of a random P-body in the existing aggregate as a destination for the new P-body. The addition of P-bodies to an aggregate follows the reaction:

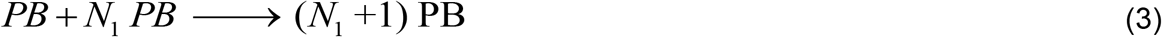

where N_1_ corresponds to the number of P-bodies in the existing aggregate. This is done iteratively until the pre-determined aggregated size is achieved. Every particle is randomly initialized with a given orientation and direction of rotation. There are six possible orientations, corresponding to the coordinate number of a triangular lattice. The direction of rotation is always clockwise (CW) or counter-clockwise (CCW). Note that, although the particle’s movement is independent of its orientation, reactant particles will only associate if their orientations are complementary.

### Statistical analysis

Graphpad-Prizm and Origin were used for statistical analysis and plotting. For pairwise comparisons, p-values were calculated based on non-parametric unpaired t-tests with Kolmogorov-Smirnov test. For comparisons involving more than 2 samples, one-way-ANOVA tests were used with Geisser-Greenhouse correction.

